# Integrated viral elements suggest the dual lifestyle of *Tetraselmis* spp. polinton-like viruses

**DOI:** 10.1101/2022.05.02.489867

**Authors:** Emily E. Chase, Christelle Desnues, Guillaume Blanc

## Abstract

In this study, we aimed at exploring horizontal gene transfer between viruses and *Chlorodendraceae* green algae (Chlorophyta) using available genomic and transcriptomic sequences for 20 algal strains. We identified a significant number of genes sharing a higher sequence similarity with viral homologues, thus signaling their possible involvement in HGTs with viruses. Further characterization showed that many of these genes were clustered in DNA regions of several tens to hundreds of kilobases in size, originally belonging to viruses related to known *Tetraselmis* spp. viruses (TetV and TsV). In contrast, the remaining candidate HGT genes were randomly dispersed in the algal genomes, more frequently transcribed and belonged to large multigene families. The presence of homologs in *Viridiplantae* suggested that these latter were more likely of algal rather than viral origin. We found a remarkable diversity Polinton- like virus (PLV) elements inserted in Tetraselmis genomes, all of which were most similar to the *Tetrasemis striata* virus TsV. The genes of PLV elements are transcriptionally inactive with the notable exception of the homologue of the TVSG_00024 gene of TsV whose function is unknown. We suggest that this gene may be involved in a sentinel process to trigger virus reactivation and excision in response to an environmental stimulus. Altogether, these results provide evidence that TsV- related viruses have a dual lifestyle, alternating between a free viral phase (i.e., virion) and a phase integrated into host genomes.

## Introduction

Polintons (also known as Mavericks) were first recognized as large transposable elements (<40Kb in size) encoding up to 10 different proteins, found in the genomes of both unicellular and multi-cellular eukaryotes (1). They generally encode both a DNA polymerase (“POL”) and a retrovirus-like integrase (“INT”), thereby how they acquired their name “POLINTons” and their classification as self-synthesizing transposons (2). Many encode a DNA packaging ATPase, major and minor jelly-roll capsid proteins (MCP and mCP respectively) and a capsid maturation protease (PRO) (3). They were therefore predicted to alternate between the transposon and viral lifestyles although virion formation remains to be demonstrated. Polintons are considered the ancient ancestors of most eukaryotic dsDNA viruses (4). There are considerable similarities between polintons and two groups of dsDNA viruses: the virophages and polinton-like viruses (PLVs). Virophages are involved in tripartite infections with giant viruses (phylum *Nucleocytoviricota)* and their cellular hosts (5, 6). Virophages can parasitize co-infecting giant DNA viruses in a host and may provide adaptive anti-giant virus defence in unicellular eukaryotes (7–9). Virophage genomes (15–30 kbp) typically encode a MCP, a mCP, a DNA packaging ATPase, and a PRO (10). They also encode DNA replication and integration proteins that were likely acquired independently in different virophage lineages (11). Some virophages can integrate into the genome of the cellular or viral host (12–14), in which case they resemble Potintons, and reactivates upon superinfection with a giant virus (8). Virophages can therefore have a dual life style combining a virus stage with an integrated stage. The PLVs are also similar in size and genomic composition to the Polintons. They have mainly been discovered in sequencing datasets and include members integrated into algal genomes (i.e. *Monoraphidium neglectum*, and *Guillardia theta*) or associated with giant virus particles (3, 15, 16). The majority of PLVs were however recovered from metagenomes of viral fractions (i.e. filtration has occurred to target particles <0.22 μm, therefore microeukaryotes have been virtually removed while viral particles remain), suggesting that PLVs occur abundantly as free viruses in the environment, especially in aquatic ecosystems (17). The PLV genomes generally do not contain all of the hallmark genes found in polintons, such as the DNA polymerase and retrovirus-like integrase, but can include ATPases, mCPs, and MCPs, and typically terminal inverted repeats at either end of their genome (3). Furthermore, PLVs were initially reported to lack the capsid maturation protease normally present in both polintons and virophages, but a recent metagenomic study suggests that a large group of PLVs contains the capsid maturation protease (17). The presence of mCPs and MCP allude to the concept of a “dual lifestyle” among PLVs as well. The first infectious PLV particle was isolated from the cosmopolitan green microalgae *Tetraselmis striata* (phylum *Chlorophyta*; family *Chlorodendraceae*) (18). The *Tetraselmis striata* virus TsV has a 31-kb genome and is estimated to release thousands of 60 nm virions per algal host cell. The TsV host infectivity range was tested on several *Tetraselmis* spp., and was only found to infect the species it was isolated from (18), highlighting the narrow host specificity of this PLV. Whether TsV also exist as an integrated form in the host genome has not yet been tested because the *T. striata* genome was not sequenced at the time of the PLV discovery. Tentative classifications of PLVs based on shared gene content and major capsid protein phylogeny, identified 3 to 5 major groups including the TSV group, which contains Tsv (3, 17). Furthermore, two other viruses have been isolated from Tetraselmis spp., including TetV, a member of Mimiviridae (“giant viruses”; phylum *Nucleocytoviricota*) with a 668-Kb genome (19). The 39Kb genome of a third potential *Tetraselmis* virus, named *Tetraselmis viridis* virus S20 (TvV-S20), is available in Genbank since 2010 and is unrelated to Tsv and TetV (20). The genetic content of this virus resembles that of an ordinary bacteriophage, suggesting that TvV-S20 could rather correspond to a bacteriophage associated with bacteria in co-culture with Tetraselmis (21).

Until recently, algal genomes of close taxonomic proximity to Tetraselmis were not available. At present, public databases contain draft genomes for several *Tetraselmis* species as well as for additional *Chlorodendraceae* genera, including *Prasinocladus* and *Platymonas* (22, 23). Transcriptomic data for several Tetraselmis strains and *Scherffelia dubia* were also made available through various sequencing programs (24–28). In this study we took the opportunity to explore both algal draft genomes and transcriptomic data for evidence of Tsv (and closely related PLVs) or other viruses potentially integrated into algal genomes (hereinafter referred to as “viral element”). More specifically our motivation was to find clues supporting the hypothesis that PLVs related to Tsv might have a dual lifestyle alternating integrated and free phases as for virophages. By searching the genomic data for viral-derived coding sequences, we identified an unexpected diversity of integrated PLV sequences closely related to Tsv in all sequenced *Chlorodendraceae* strains. In addition, we highlight large genome fragments of giant TetV-related viruses in the *T. striata* and *T. suecica* genomes with remarkably different GC contents than the original TetV.

## Materials and Methods

### Acquisition of genomic and transcriptomic data from family Chlorodendraceae

Six draft genome assemblies for *T. striata* (strains LANL1001 and UTEXSP22), *T. suecica* (strain CCMP904 sequenced independently by New York University Abu Dhabi [NYUAD] and Institut Français de Recherche pour l’Exploitation de la MER [IFREMER]), *T. subcordiformis* and *Prasinocladus* sp. malaysianus were retrieved from the National Center for Biotechnology Information (NCBI) GenBank, the Dryad database and the IFREMER data repository server. Genome sequencing of *T. striata* LANL1001 and *T. suecica* CCMP904 (IFREMER version) was performed using long read sequencing technologies (PacBio and ONT, respetively), which resulted in longer assembled contigs. Also, transriptomic data (Illumina RNA-seq reads) for 15 Chlorodendraceae strains, including *Scherffelia dubia* and several *Tetraselmis* spp., were retrieved from Genbank. Algal strain names and identifying information for retrieving sequence data are given in **Table S1**. Genbank accession numbers of the virus genomic sequences used in this study are KY322437 for TetV, NC_020869 for Tsv, and NC_020840 for TvV- S20.

Note that the *T. suecica* CCMP904 assemblies released by NYUAD and IFREMER were quite dissimilar. The NUYAD assembly was generated using Illumina short reads, and contained a large number of contigs (N=392,918) of relatively short length (<20Kb, L50=868bp), totaling 237 Mb (23). In contrast, the IFREMER assembly was generated using Oxford Nanopore Technologies (ONT) long reads, and contained a much smaller number of contigs (N=175) of greater sizes (L50=4.3Mb), totaling 100 Mb. Mapping of 39 million NYUAD short reads on the IFREMER assembly using HISAT2 (29) resulted in 30% of overall alignment rate. Likewise, 27% of the NYUAD assembly sequence found a significant BLASTN match (e-value <1e-100) in the IFREMER assembly. Conversely 65% of the IFREMER assembly sequence was aligned on the NYUAD assembly. Thus the two assemblies had only partial overlap and contained a substantial amount of assembly-specific sequences. Part of the unmatched sequences is the consequence of the high sequencing error rate induced by the ONT technology (i.e., the overlap between the two assemblies is therefore underestimated).

### Sequence analysis

Transcriptome raw reads were first trimmed for low-quality and adaptor sequences using AlienTrimmer (30), then assembled into transcript sequences using SPADES v3.15.3 (31) with the flag “--rna”and default parameters otherwise. Multiple RNA-seq datasets from the same strain were pooled before assembly. Finally, reconstructed transcripts were clustered based on nucleotide similarity (>95%) using CDHIT-EST (32) to remove redundancy due to alternative splicing and highly conserved paralogues. The longest sequence of each sequence cluster was kept for further analysis.

For each assembly (transcriptome or genome), we generated a collection of all open reading frames >150 codons (see **Table S2** for assembly and ORFing statistics). While the typical structure of eukaryotic algal genes contains introns, the use of an ORFing approach (i.e. without consideration of introns that interrupt the reading phase) in the present genomes is not a problem because viral genes do not have introns, which is generally true for genes brought by viral genome insertions. Translated ORFs were aligned against the TrEMBL public database using MMSEQS (e-value <1e-5) (33) and the best matches were recorded. We did not consider best matches to uncultured organism or environmental sequence because the nature of the corresponding organism is often uncertain. We also disregarded matches with *Chlorodendraceae* species to allow detection of putative viral HGT events that occurred in a *Chlorodendraceae* ancestor. When a translated ORF returned a best match to one of these sequences, we recorded the next highest scoring match to a clearly identified non-*Chlorodendraceae* organism or virus. Finally, ORFs with a recorded best match to a virus (“VBM” will be used hereafter in place of “viral best match”) were considered as potentially involved in HGT with a virus and were analyzed further. Note that this approach may miss some candidates when the best match is a viral insertion in the genome of a cellular organism. Protein family clustering was done using the MMSEQS easy-cluster workflow by setting the -c parameter to 0.2 to account for truncated sequences.

Transcriptomic analysis was performed using the HISAT2 program (29) with default parameters to map RNA-seq reads onto the reference genomes. The resulting alignments were used to calculate the FPKM values (Fragments Per Kilobase Million) for gene, which are used as an estimation of the relative transcription level in the cell.

### Identification of putative viral elements based on VBM identification

Viral elements were defined as clusters of candidate viral genes co-localized on a genomic contig or segment of thereof. We applied two types of methods for detecting these clusters. The sliding-window approach consisted of sliding a 10-Kb window along the contig with a 1-Kb step and counting the number of VBM ORFs in each window. We systematically investigated genomic regions returning windows that had at least four candidate viral genes. This approach allowed to identify large viral inserts of TetV-like DNA. In the second approach, which proved to be more effective in identifying conserved PLV elements (shorter insertions), the MCP of Tsv was used as “bait” to recover all potential MCPs from each of the algal genomes. Then, 20 Kb upstream and downstream of algal MCP gene were searched for additional ORFs with a best match to the Tsv genome or to a collection of PLVs genomes gathered in Yutin et al. 2015 (3). Finally, sequences of 3 Kb following the last matches against the PLVs on the left and on the right were aligned by BLASTN to identify possible direct or reverse repeats (i.e. terminal inverted repeats; TIRs) likely to mark the limits of the PLV elements.

### Phylogenetic placement of relevant algal strains by RBCL and VBMs by MCP genes

A phylogenetic tree of the algal strains used in this study was reconstructed based on the RBCL coding sequence (large subunit of ribulose bisphosphate carboxylase) identified in Genbank NR and in the assembled datasets using BLAST (**Figure S1**). A multiple sequence alignment of the RBCL genes was generated with MAFFT (34), followed by a maximum likelihood phylogenetic reconstruction using the FastTree program (35) and the GTR+G substitution model. Statistical support for branches was estimated using the Shimodaira-Hasegawa test as implemented in FastTree. The RBCL tree showed 2 apparent incongruences between species names and phylogenetic position. The first one concerns *T. suecica* PLY305 which clusters within the *T. striata* clade while other algae referenced under the *T. suecica* species name grouped in their own clade. The corresponding transcriptomic sequences came from the oneKp project (25), a large-scale sequencing program, and it is possible that there has been a confusion in the identification of the sequenced material. There was also a fairly large phylogenetic distance between *T. subcordiformis* FACHB-1751 and UTEX171, which suggests that these two organisms did not belong to the same species. The phylogenetic trees based on protein sequences were produced using the same procedure except that we used the WAG+G substitution model adapted to protein sequences.

## Results

### Virus-like genes recovered from algal genomes of family Chlorodendraceae

ORFs with a viral best match (VBM) in TrEMBL were identified in the genome assemblies of *T. striata, T*.*suecica, P. subcordiformis*, and *Prasinocladus* sp. malaysianus. The number of these ORFs ranged from 51 for *Prasinocladus* sp. *malaysianus*, to 1148 for *T. striata* LANL1001. Overall VBMs were primarily from proteins of the Tetraselmis infecting viruses including Tsv and TetV (see **Figure 1A**). However, we obtained no best hits to TvV-S20 for all tested species. A smaller fraction of VBMs were from members of families *Phycodnaviridae, Mimiviridae*, and *Lavidaviridae* (i.e. virophages) – of which the first two groups contain microalgae infecting viruses and are giant viruses (phylum *Nucleocytoviricota*). Additionally, there were few VBMs to other known giant viruses (i.e. families Irodoviridae, Marseilleviridae, and Pitoviridae). Overall, over 80% of all hits were from Tsv or TetV, with TetV responsible for 58% of the total. The proportion of Tsv and TetV hits varied across species and also across strains of the same species, with noticeable differences between both *T. striata* strains (Tsv was 29% and 41%, and TetV was 52% and 22%, for LANL1001 and UTEXSP22 respectively) and between *T. suecica* CCMP904 assemblies (Tsv was 10% and 17%, and TetV was 71% and 53% for NYUAD and IFREMER, respectively). Overall, the *T. suecica* CCMP904 NYUAD assembly had 4 times more VBM than the *T. suecica* CCMP904 IFREMER assembly. These incongruences may have different reasons, including the heterogeneous quality of genome assembly, sequencing depth and technology, and different within-species history of interaction with viruses. Because of these potential technological biases, VBM gene counts in CCMP904 and other algae should be taken with some caution. Although *Prasinocladus* sp. malaysianus diverged early from the *Tetraselmis* genus within the *Chlorodendraceae* family (**Figure S1)** it also had viral hits (albeit substantially less than other algal genomes analysed), which included both Tsv and TetV matches; two viruses not known to be associated with genus *Prasinocladus*.

**Figure 1.**
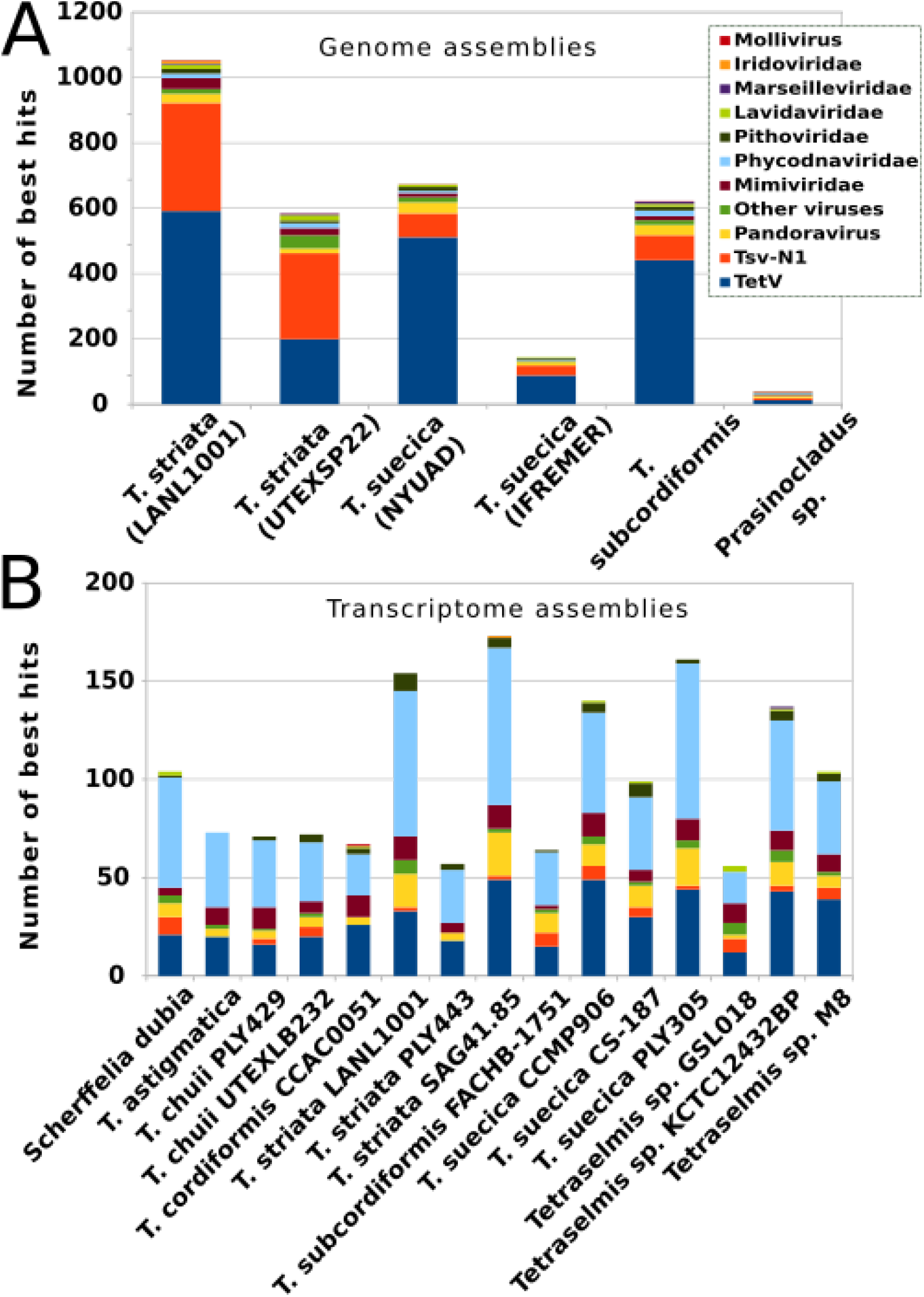
Taxonomic classification of TrEMBL viral best matches from *Chlorodendraceae* derived sequences. **(A)** Number of corresponding best matches in ORFs generated from draft genomes of several *Tetraselmis* spp., and a member of genus *Prasinocladus*. **(B)** Viral best matches in *Chlorodendraceae* transcripts. Colours correspond to the best match’s family, except in the case of Tsv (a polinton-like virus), and TetV (member of family *Mimiviridae*).

To complete the analyses we assembled public transcriptomic data from 15 cultured *Chlorodendraceae* and searched for VBM transcripts. VBM transcripts were found in all transcriptomes (**Figure 1B**), but their counts were generally lower than in algal genome assemblies. *T. striata* LANL1001 was the only strain for which both genomic and transcriptomic data were available. Considerably more VBM genes were identified in the genomic assembly than in the transcriptomic assembly, i.e., 1148 VBM ORFs versus 149 VBM transcripts, respectively. This suggests that only a fraction of the LANL1001 VBM ORFs were transcribed under the culture conditions applied for transcriptome sequencing. Furthermore, the taxonomic distribution of VBMs differed substantially between ORFs and transcripts as shown in **Figures 1A and 1B**, respectively. *Phycodnaviridae* was the dominant taxonomic category in transcript VBMs, recruiting from 31% to 58% of VBMs, whereas it only accounted for 6 to 18% of the ORF VBMs. TetV also recruited a large portion of the transcript VBMs (21-41%) but less so than for genomic ORFs (31-71%). Finally, Tsv accounted for 0-10% of transcript VBMs, which is substantially less than the 10-41% observed in genomic ORFs. Overall, these results suggest that the propensity of a gene to be transcribed varies with the taxonomic origin of its VBM under standard culture conditions.

Algal sequences with a VBM (3431 genomic ORFs and 1532 transcripts; **Dataset S1**) were translated and clustered into 561 protein families (**Table S3**). About two thirds of the protein families (N=344) were found in two or more strains. The largest family included 802 proteins containing a RING-type z-finger domain (FAM001). These proteins were encoded by highly duplicated genes in all *Chlorodendraceae*. Only one remote homolog was found in green algae (i.e., Trebouxia sp. A1-2 KAA6422835). Their most closely related homologs were found in Chloroviruses which contained from 1 to 10 gene copies. Other large protein families include Ankyrin domain proteins (FAM002, N=367) also found in various Nucleocytoviricota, homologues of the TsV (PLV) DNA packaging ATPases (FAM003; N=236) and homologues of the TsV MCP (FAM004; N=200).

We queried the *Chlorodendraceae* sequences with reference major capsid proteins (MCP) from TetV (Nucleocytoviricota) and from TetV (PLV). This approach is more sensitive than the former VBM analysis because some algal sequences may be truncated or match viral insertions in cellular genomes, in which case they cannot be detected as VBM candidate. By definition, MCPs are encoded by hallmark viral genes and therefore cannot be of cellular origin; their presence necessarily attests to events of genetic transfer from viruses to algae. The amino acid sequence divergence between Nucleocytoviricota MCPs and PLV MCPs is generally so high that they cannot align against each other at the specified threshold of detection (MMSEQS e-value<1e- 5). In addition to the number of matching ORFs in genome assemblies, Table 1 reports the number of contigs carrying these ORFs, because sequencing errors or nucleotide substitutions can lead to splitting of a full length gene into several successive ORFs (hence multiple counts) due to occurrence of internal stop codons. This contig number can be considered as a lower bound on the estimate of the number of actual MCP genes in each genome assembly. Thus, between zero and four ORFs had a significant protein similarity to the TetV MCP and between two and 247 ORFs (on 203 contigs) had significant protein similarity to the TsV MCP (**Table 1**). In transcriptomes, only *Tetraselmis* sp. KCTC12432BP had a transcript with significant similarity to the TetV MCP, and 8 distinct Chlorodendraceae strains had between 2 to 10 transcripts with significant similarity to a PLV MCP.

**Table 1.** Total number of best matches to the virus hallmark MCP

|  | **TetV MCP\*** | **TsV MCP\*** |
| --- | --- | --- |
| **Genome assemblies:** |  |  |
| *T. striata* LANL1001 | 1 (1) | 126 (98) |
| *T. striata* UTEXSP22 | 1 (1) | 247 (203) |
| *T. suecica* CCMP904 NYUAD | 4 (4) | 105 (100) |
| *T. suecica* CCMP904 IFREMER | 0 | 39 (9) |
| *T. subcordiformis* UTEX171 | 3 (3) | 115 (103) |
| *Prasinocladus* sp. malaysianus UTEX2601 | 0 | 2 (2) |
| **Transcriptome assemblies\*\*:** |  |  |
| *Scherffelia dubia* CCAC0019 | 0 | 3 |
| *T. chuii* PLY429 | 0 | 2 |
| *T. chuii* UTEXLB232 | 0 | 4 |
| *T. subcordiformis* FACHB-1751 | 0 | 2 |
| *T. suecica* CCMP906 | 0 | 10 |
| *T. suecica* CS-187 | 0 | 9 |
| *Tetraselmis* sp. GSL018 | 0 | 7 |
| *Tetraselmis* sp. KCTC12432BP | 1 | 0 |
| *Tetraselmis* sp. M8 | 0 | 4 |

\* number of ORFs (number of contigs); \*\* transcriptome datasets are listed only if they contained a best match to one of the two target MCPs.

Phylogenetic reconstruction of the TetV-like MCPs indicated that most algal sequences clustered with the TetV (Tetraselmis giant virus) homologue, further confirming the close-knit relationship between this viral clade and the *Tetraselmis* genus (Figure 2). However, the MCP encoded by the *Tetraselmis* sp. KCTC12432BP transcript was more divergent and occupied an isolated position in the tree without close viral homologues. This sequence could represent the signature of a still unknown virus infecting this Tetraselmis strain. A phylogenetic analysis of the PLV-like MCP is described in the PLV elements section below.

**Figure 2.**
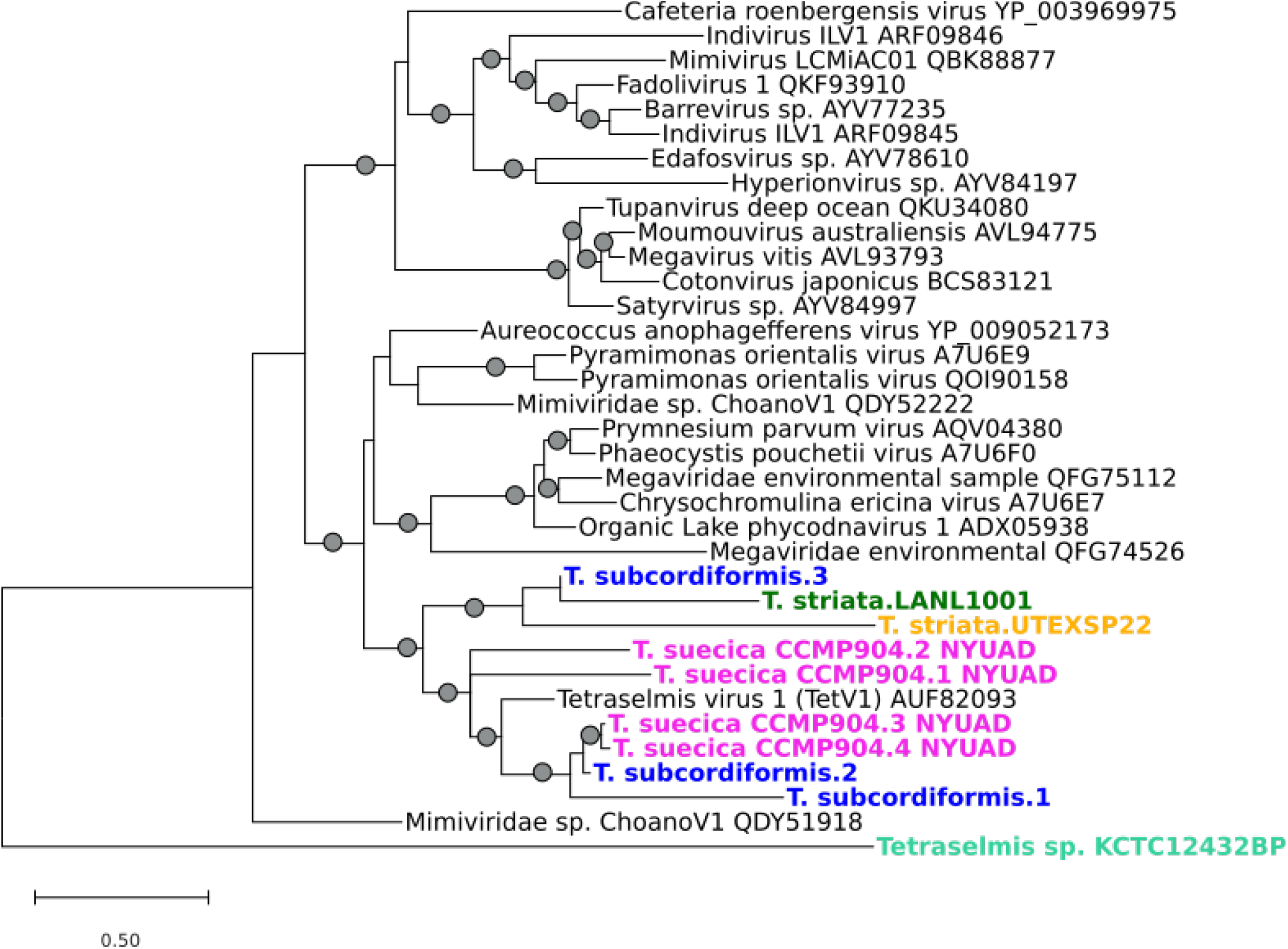
Unrooted phylogenetic tree of major capsid proteins (MCP). Proteins identified in the *Chlorodendraceae* data are indicated in color. Grey circles show branches with >85% local support as calculated with the Shimodaira-Hasegawa test. The branch scale indicates the number of amino-acid substitution per site.

### Large TetV-like genome insertions in Chlorodendraceae algae

We then analysed the distribution of VBM genes in the genome assemblies using a sliding window approach. Contigs containing 10-Kb windows with at least four viral ORFs were identified in all assemblies but *Prasinocladus* sp. and were inspected further using graphical representations of the ORF best hit taxonomic classifications and local GC content of contigs. This way we could identify three large regions (roughly 400Kb to 500Kb) of the *T. striata* LANL1001 assembly that had a higher viral gene density (**Figure 3**). We confirmed that the regions were flanked by ORFs with algal matches to rule out viral contamination. Using a phylogenetic analysis, the relationship of the integrated viral genome segments to TetV was confirmed using the single copy of DNA polymerase ORF identified in each fragment. The TetV-like MCP gene identified in LANL1001 (**Table 1**) is not located in one of these genomic regions; instead, it is isolated among genes of algal origin suggesting that it results from another integration event. In regard to the host GC content verses the viral insertions, one located on contig #2972 is more similar (G+C=58.8%) to the GC content of the host algal genome (G+C=58.0%), whereas the two others are drastically different at over 66% GC content (G+C=66.0% and 67.1% for contigs #3242 and #784 respectively) (**Figure 3**). Furthermore, the CG content of the viral genome of TetV is also drastically different (i.e. 41.2%) than that of all three viral insertions and the host algal genome.

**Figure 3.**
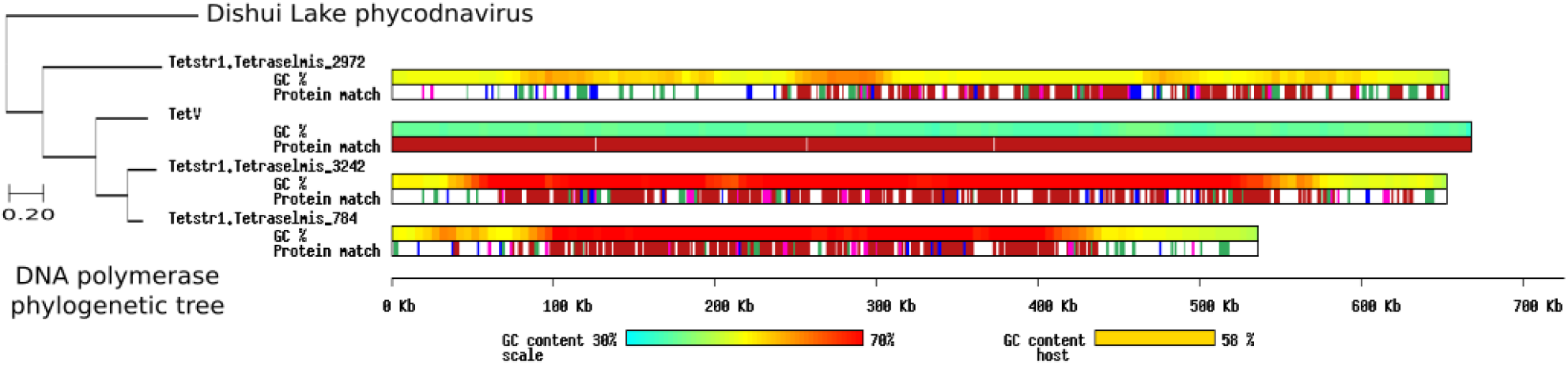
Comparison of three viral insertion elements within a *Tetraselmis striata* draft genome (LANL1001), with the corresponding viral match; TetV-1. ORFs are represented in the “protein match” track, where red represent best match in viruses, green are for best match in green algae (i.e., Chlorophyta), blue are for best match in eukaryotes and pink represent best match in prokaryotes. GC content is also reported for each viral elements/TetV, alongside the GC content of the host genome (*T. striata*) overall. On the left, a phylogenetic tree is shown based on viral DNA polymerase identified as a single copy ORF in each contig and virus. The phylogenetic tree is rooted by the Dishui Lake phycodnavirus homologue. The scale bar represents the number of amino acid substitution per site.

Although other investigated algal genomes contained a great number of best matches to TetV (i.e. *T. striata* UTEXSP22, *T. suecica* CCMP904 NYUAD and *T. subcordiformis* UTEX171), we did not identify such large viral insertions in their assemblies (see **Figure S2**). One obvious reason is that these Illumina short read assemblies were highly truncated as reflected by the quasi absence of contigs >20 Kb, which was not suitable for this analysis. As a result, positive windows were only located on short contigs, which did not contain additional ORFs matching to green algae. Thus, we could not determine if these sequences originated from viral contaminants or integrated viral DNA. All of the contigs containing positive windows shown in **Figure S2** had a majority of best matches to TetV. Their G+C contents ranged between 47.2% and 63.9%, and were generally higher than the average host G+C% (from 51.6% for *T. suecica* CCMP904 NYUAD to 56.4% for *T. striata* UTEXSP22).

In contrast, *T. suecica* CCMP904 IFREMER project was sequenced using long ONT reads, and 97% of the genomic sequences was assembled in chromosome-sized contigs >100 Kb. Although the number of TetV matches for this genome assembly is comparatively low, we found three contigs of likely viral origin, generating positive windows containing exclusively best matches to TetV. The largest contig (#6216) was 383 Kb with a GC content (63.6%) higher than for the algal (53.0%) and TetV genomes. The two remainders (#5310 and #5311) were 11 Kb and 32 Kb (72.5% and 73.9% G+C, respectively). Five additional high GC content contigs (G+C>70%; contigs #6197, #6058, #1543, #2055 and #5034) containing best matches to Nucleocytoviricota (including TetV) but too short to generate positive windows were also observed in the graphical representation of VBM genes in the *T. suecica* CCMP904 IFREMER genome assembly (**Figure S3**). All of these high-GC contigs could originate from the same genomic region. Again, the absence of match to green algae at the extremity of the contigs made it impossible to determine if these sequences result from viral contaminants or integrated viral DNA. An analysis of the depth of coverage of IFREMER contigs by Illumina NYUAD reads indicated that algal contigs have an average mapping coverage of 43X whereas viral contigs have an average coverage of 3.3X (**Figure S4**). In the case of virus genome fragments integrated in single copy in the algal genome, an equal coverage should make most sense. Two alternative hypotheses can explain this incongruence: (i) DNA fragments with the most extreme GC contents are generally underrepresented in sequencing results with Illumina technology (36). Thus the lower coverage in viral contigs, which have a substancially higher GC content than the algal contigs, could be a simple manifestation of this systematic bias. (ii) It is also possible that the viral contigs are not part of the algal genome but rather originate from a low intensity viral infection contained in the CCMP904 cultures distributed by the culture collection centers. Other genes with a best match in *Nucleocytoviricota* but located in genomic regions with a background GC content appear to be randomly distributed. The largest TetV-like *T. suecica* CCMP904 IFREMER contig and the 3 *T. striata* LANL1001 contigs exhibited residual gene colinearity conservation with the TetV genome and with each other (**Figure S5**).

In all cases, our results suggest that five strains of Tetraselmis, which are distinct from the Tetraselmis strain on which TetV was initially isolated (19), were concerned by infection by TetV relatives at some point in the history of their lineages. Collectively, the algal genomes contained copies for 285 of the 653 protein genes predicted in the TetV genome, with *T. suecica* CCMP904 IFREMER having the smallest TetV gene complement (N=51) and *T. striata* LANL1001 the largest (N=253).

### PLV-like elements provide insights into the evolution and mechanisms behind PLVs

A total of 66 integrated viral elements above 2000 bp in length and with similarities to PLVs were also retrieved from the six draft genomes assessed in this study. We also identified 16 contigs resembling PLVs in the assembled transcriptomic data of 7 *Chlorodendraceae* strains including *Tetraselmis* spp. GSL018 and M8, *T. subcordiformis, T. chui* UTEXLB232, *S. dubia, T. suecica* CS-187 and CCMP906. These contigs often contained multiple ORFs, sometimes in opposite directions and/or corresponding to distinct hallmark PLV genes, which probably reflects sequence overlaps between successive PLV transcripts. Phylogenetic reconstruction based on the PLV MCP shows a remarkable diversity of the algal PLV-like elements around the TsV itself (**Figure 4A**). Four coherent phylogenetic groups could be defined (G1 to G4), one of which contains the TsV (G2). The closest TsV relative was a short transcribed fragment assembled from the *T. subcordiformis* FACHB-1751 RNA-seq data (76% amino acid identity between MCPs). Elements from the same algal species, such as *T. striata* or *T. suecica*, tended to share higher sequence similarity and clustered tightly in the tree. This suggests that algal species independently underwent burst of insertions by specific viral lineages rather than steady accumulation of insertions by a diversity of PLVs. Structural annotation revealed many repeating viral gene hits across the PLV-like elements, but their relative order varied especially between phylogenetic groups (**Figure 4B**). Several genes are found split into several consecutive ORFs due to interior stop codons in the reading phase, following sequencing errors or sequence decay. Shared genes included ones from a “typical” PLV repertoire and encode proteins such as ATPase, minor and major capsid proteins, as well as genes noted by previous studies of PLVs (3) including lipase, cytosine methylase and transcription factor VLTF3. Lipases are enzymes that normally break down lipids but have been recorded without this activity in viruses and instead could be involved in replication (37). Four shared putative genes without a known or implied function (TVSGs and G-protein (3); i.e. hypothetical proteins) were also recorded and of significance because they appear in TsV and other PLVs. Interestingly PLV-like elements had 3 different, mutually exclusive genes encoding proteins involved in viral genome replication, including S3H helicase – TVpol (transposon-viral polymerase) fusion, protein-primed DNA polymerase (pDNAP) and S1H helicase. This suggests that these important genes can be readily interchanged during PLV evolution but they seemingly cannot coexist in a same viral genome. The most parsimonious scenario based on conciliation of gene distribution and tree topology suggests that S3H helicase – TVpol was the ancestral gene form and was replaced by the S1H helicase in group G2 PLV-like elements and pDNAP in group G4. However phylogenetic reconstructions of these proteins show complex evolutionary trajectories. Although the pDNAP-bearing elements cluster within the G4 group according to the MCP phylogeny, the phylogenetic tree of pDNAPs suggests that they originated in two distinct phylogenetic groups (i.e., 1 and 2 in green in Fig. 4 and Fig. S6A). This suggests repeated colonizations of *Chlorodendraceae* PLVs by pDNAP genes of different origins during recent evolutionary history. The closest relatives of the *Chlorodendraceae* pDNAP are found in eukaryotic genomes, possibly encoded by ancient PLV elements that invaded those genomes. The phylogenetic reconstruction of S3H helicase - TVpol fusion proteins sends a similar message. The *Chlorodendraceae* PLVs carrying this gene belong to the G1, G3 and G4 groups defined by the MCPs phylogeny. However, within the G1 group, the S3H helicase - Tvpol genes are from two distinct phylogenetic groups (i.e., 1 and 2 in orange in Fig. 4 and Fig. S6B), again suggesting repeated colonizations by genes of different origins. Again, the closest relatives are mainly found in other eukaryotic genomes. The S1H proteins of PLVs did not show sufficient sequence similarity to other homologs to allow robust phylogenetic reconstruction. Thus their origin could not be established.

**Figure 4.**
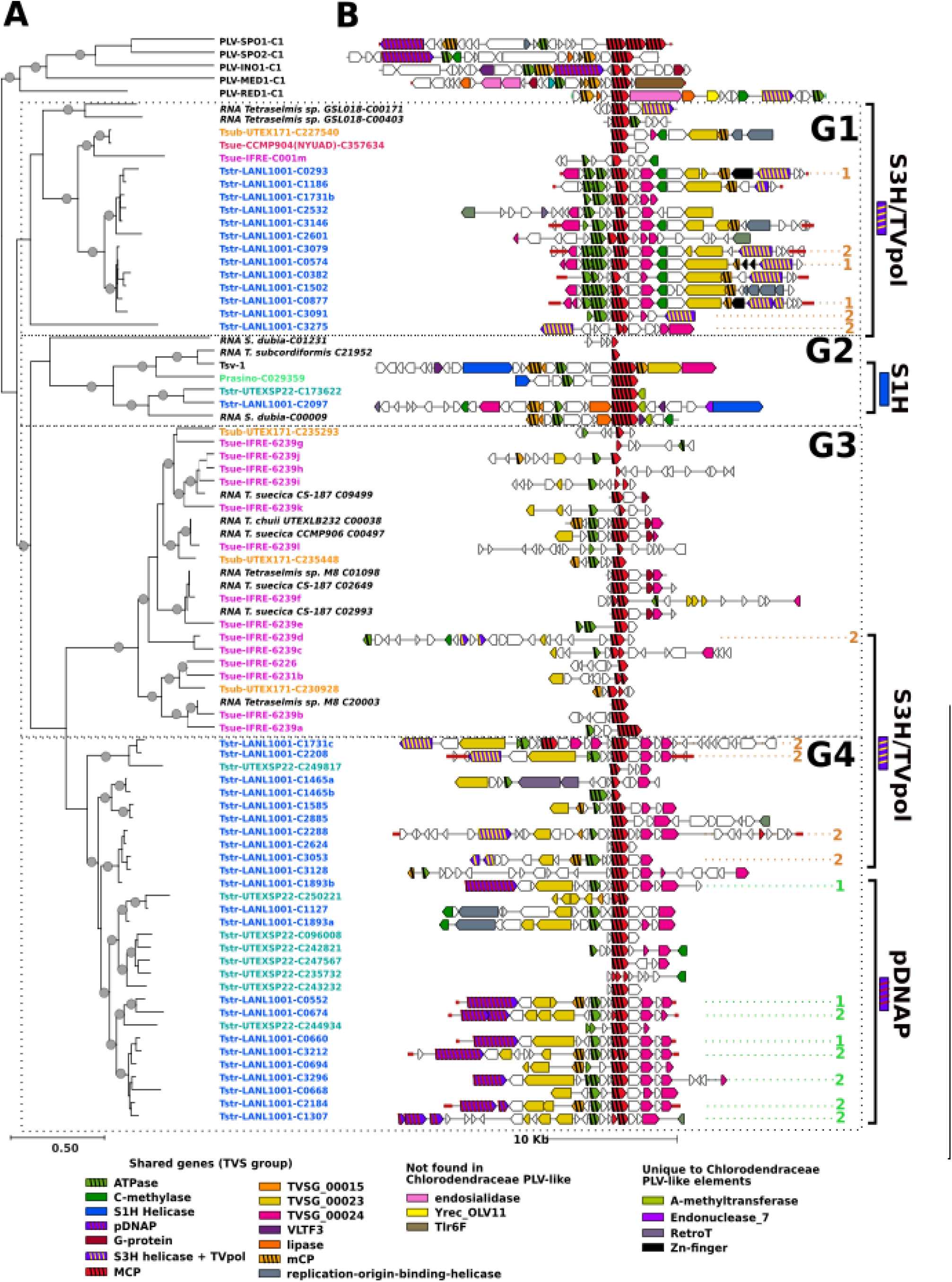
PLV-like elements in algal genomes. **(A)** Phylogenetic tree of the MCPs encoded by PLV-like elements, rooted by several PLVs previously identified as members of the TVS group by Yutin et al. (3). **(B)** Genomic architecture of the corresponding PLV-like elements. Names of elements identified in genomic data are shown in color whereas names of elements identified in transcriptomic data are in black and preceded by the term “RNA”. We only considered PLV elements >2000 bp except RNA *T. subcordiformis* C21952 (565 bp) which was conserved owing to its close phylogenetic proximity with TsV. Structural annotations of each viral elements are depicted with a colour/pattern legend. Spans of terminal inverted repeats (TIRs) are identified on relevant viral elements by red lines. Scale of the phylogenetic tree (amino acid substitutions/site) and scale of the genome lengths are included. Abbreviations are RED: Red Sea, MED: Mediterranean, INO: Indian Ocean and SPO: South Pacific Ocean, according in reference to their sampling locations; IFRE, IFREMER. Tstr: *T. striata*; Tsue: *T. suecica*; Tsub: *T. subcordiformis*; Prasino: *Prasinocladus* sp. *malaysianus*. Transcribed PLV-like sequences are shown in italic. Numbers (1 and 2) in green and orange indicate the clustering of the pDNAP and S3H/Tvpol proteins in their respective phylogenetic trees shown in Fig. S6A and B, respectively.

No significant hit to protein involved in DNA integration (e.g., RVE integrase or tyrosine recombinase YREC (OLV11)) was found in *Chlorodendraceae* sequences, either within or outside the identified PLV-like elements. Thus the mechanism and enzymatic apparatus responsible for the *Chlorodendraceae* PLV integration remain unknown. Other genes previously found in PLVs of the TVS group but missing from the *Chlorodendraceae* elements are the ones encoding Tlr6F and endosialidase. Tlr6F is an uncharacterized protein found in several viruses (e.g. PLVs, *Lavidaviridae, Nucleocytoviricota*, and bacteriophages) and thought to be a component of viral reproduction (3).

Some of the *Chlorodendraceae* PLV element genes have not been documented in PLVs of the previoulsy described Tsv group yet (3). For example zinc finger (displayed as Zn- finger) proteins encoded by some group G1 PLV elements are known to have many different functions in relation to interacting with DNA or RNA, or other proteins, but in some cases have been shown to inhibit replication in viruses such as retroviruses, alphaviruses, and filoviruses. These are RNA virus taxa, however, designed zinc finger proteins have been used to inhibit a clinical DNA virus (herpes simplex virus 1 (38)). Zinc finger domains have also been shown to be associated with an integrase domain in eukaryotic transposons (39). It is possible that these zinc finger genes could have some effect on replication or have been involved in the PLV integration process. PLV elements of group G2 carry a gene encoding an adenine-methyltransferase of prokaryotic origin. This gene has never been identified in other PLVs of the previoulsy described Tsv group (3) but distant homologues are found in Yellowstone lake phycodnavirus 2 and in some bacteriophages. In the latter, adenine methyltransferase activity renders the DNA resistant to cleavage by restriction enzymes sensitive to adenine methylation. Moreover, these same PLV elements carry in the vicinity of the adenine-methyltransferase gene another gene coding for a restriction endonuclease also of prokaryotic origin. It seems that these PLVs have a complete restriction- modification (RM) system that probably allows them to recognize and degrade the DNA of unmethylated competitor viruses in case of coinfection. Methyltransferases and putative RM systems have been domumented in about half of the PLVs and virophages detected in some recent metagenome studies (17, 40) highlighting a likely importance of these functions in the replicative cycle of these viruses. Finally, several PLV elements of the G2 and G4 groups carry a retrotransposon. It is possible that these transposable elements were inserted into the PLV elements after the latter were integrated into the genome. However, the position of the retrotransposon is conserved in PLV elements of the same group, suggesting that the insertion of the retrotransposon occurred in the ancestor of these PLVs, before the latter integrated into the host genome. If this hypothesis is true, the PLVs could then participate in the propagation of transposable elements from one host to another. A previous genomic study of the marine heterotrophic protist *Cafeteria burkhardae* also revealed a close association between integrated virophages and a family of retrotransposons (12), highlighting again the potential role of viruses in horizontal transmission.

Some PLV-like elements exhibited terminal inverted repeats (TIRs) at both ends of the sequence (**Figure 4B)**. Typically TIRs indicate that the genome of the inserted PLV is complete (3, 41, 42). There is a total of 13 PLVs retrieved from these datasets that have TIRs flanking the genes (i.e. on both ends), suggesting that these 13 PLV insertions are likely complete. Their sizes ranged from 16,954 bp (Tstr-LANL1001- C0660) to 31,706 bp (Tstr-LANL1001-C2288), with a mean at 19,903bp, which is smaller than the size of TsV (26,407 bp) but comparable to other PLVs (MED1: 21,217bp, RED1: 19,641 bp, SPO1: 22,704 bp (3)). Another interesting feature is the apparent uneven distribution of PLV-like elements along chromosomes of *T. suecica* CCMP904 IFREMER (**Figure S3**). 56% of the ORFs encoding a PLV MCP, including group 1 and 2 MCPs (**Figure 4**), were clustered in the central region (between positions 8,6 Mb and 9.4 Mb) of the largest contig (Tsue-IFRE-6239; 17,2 Mb), which most likely correspond to the largest chromosome. The remaining PLV MCP genes were randomly dispersed throughout the rest of the genome. This central region may represent a hotspot for PLV integration. Unfortunately the highly fragmented assemblies of the other algal species made it impossible to determine whether such hotspots also exist in other *Chlorodendraceae*.

### VBM gene transcription were low among TsV and TetV categories within Chlorodendraceae individuals

We sought to study the transcription activity of VBM genes using the available *Chlorodendraceae* RNA-seq data. Only *T. striata* LANL1001 had both genomic and RNA- seq data available to perform a conventional transcriptomic analysis (i.e., genome and transcriptome from the same strain). To widen the analysis, we also aligned RNA-seq data and genome assemblies from the same phylogenetic group (i.e., the *T. striata* and *T. suecica* groups as defined in the RBCL phylogenetic tree as shown in **Figure S1**), assuming that VBM genes are likely to be conserved between closely related strains. Alignment of the *T. striata* LANL1001 transcriptomic data on the *T. striata* LANL1001 genome assembly revealed that most VBM genes associated with the taxonomic categories TetV and TsV were transcriptionally silent (FPKM∼0; see **Figure 5A and Tables S4-S5**). This was also the case for VBM genes associated with the category “Other viruses”, however these genes were most often located within PLV elements or in TetV-like genomic regions, suggesting that they have the same evolutionary origin as the TsV or TetV categories. Only 16, 26 and 2 genes from categories “TsV”, “TetV” and “Other” had FPKM≥1 and were considered as transcribed (accounting for 2.7%, 7.9% and 3.9% of their categories, respectively). Being outliers relative to the bulk of the genes that are non-transcribed in each category, they are not apparent in the boxplots in Figure 5A. Interestingly 13 of the 16 transcribed TsV-like genes were most similar to the unknown protein gene TVSG_00024 (found extensively in G1 and G4; **Figure 4B**), suggesting that they may act as sentinel genes that orchestrate a possible reactivation of virus following an unknown stimulus or event. The protein shares partial similarity with bacterial homologs within a region containing a discoidin domain. Discoidin domains are found in a variety of eukaryotic and prokaryotic proteins characterized by a considerable functional diversity (43). Transcribed genes from the “TetV” and “Other” categories had more diverse functions. RNA-seq data of four other Tetraselmis strains did not align better with the LANL1001 VBM genes in categories “TsV”, “TetV” and “Other”. In contrast, a large fraction of VBM genes in categories Mimiviridae, Pandoraviruses, Phycodnaviridae and Pithoviridae were transcribed in all five Tetraselmis strains (**Figure S7**). The same analysis using the *T. striata* UTEXSP22 genome as reference yielded similar overall results (**Figure 5B**). A different set of RNA-seq data was aligned against the genome assemblies of the suecica group. When the *T. suecica* CCMP904 IFREMER genome was used as reference, the measured levels of transcription were generally null or very low for genes in the TsV, TetV and “Other” categories in all tested transcriptomes whereas they were more substantial for genes in the Mimiviridae, Pandoraviruses, Phycodnaviridae and Pithoviridae categories (**Figure 5C**). This overall pattern was similar to what was observed in the striata group. In contrast, when the genomes of *T. suecica* CCMP904 NYUAD and *T. subcordiformis* UTEX171 were used as reference, only the *Phycodnaviridae* VBM gene category exhibited substantial gene transcription (**Figures 5D and 5F**). The difference observed between the IFREMER distribution and that of the other 2 strains of the suecica clade, especially for the categories Mimiviridae, Pandoravirus and Pithoviridae, is mainly explained by the fact that the IFREMER strain contained comparatively small numbers of genes in these categories (**Table S5 and Figure S7**). These genes were also mostly transcribed. They also exist in the other two strains of the suecica group, and are also transcribed, but are outnumbered in their category by a large number of additional genes that are not transcribed.

**Figure 5.**
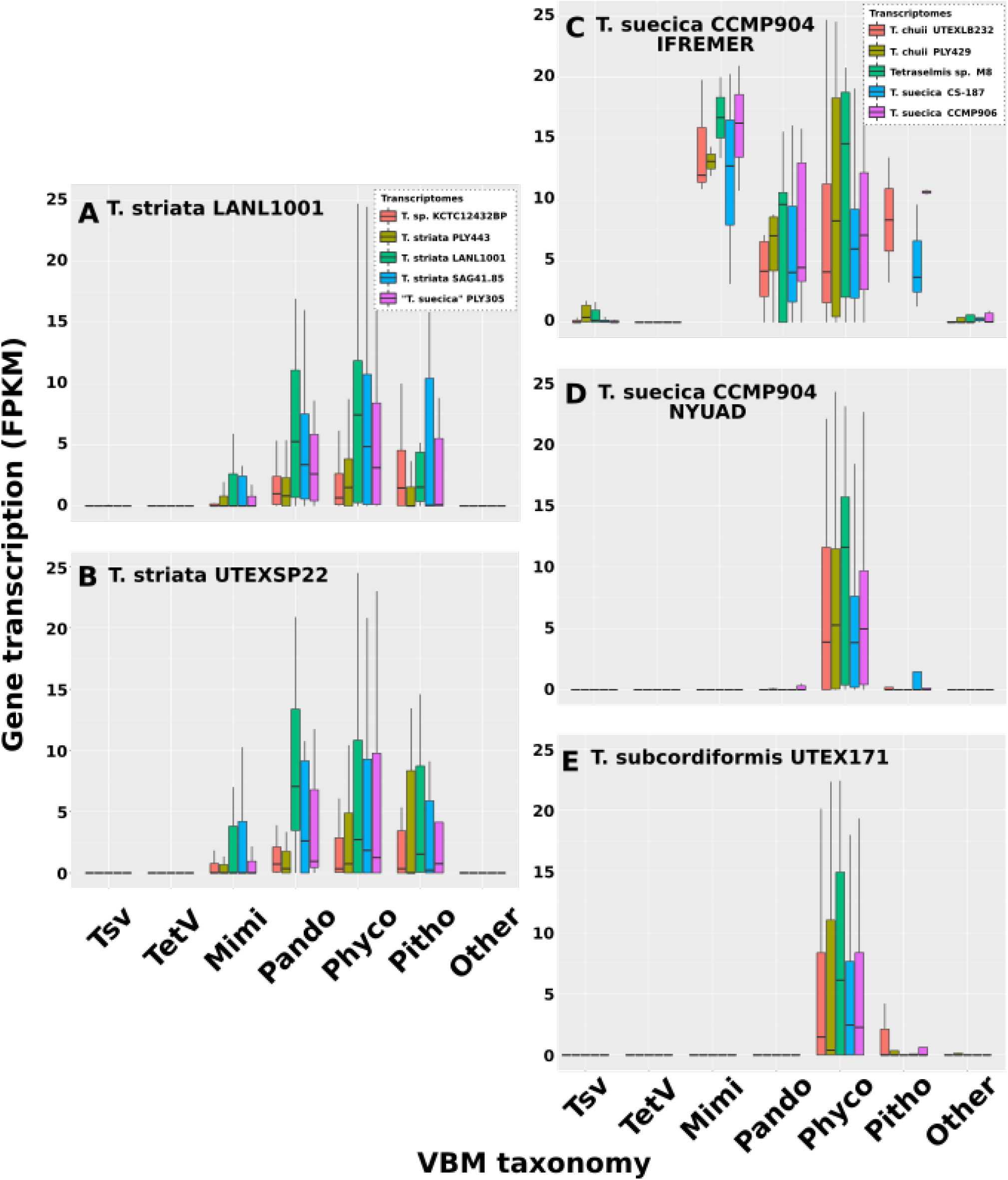
Transcription activity of VBM genes. Five algal genomes **(A—E)** were used as reference to map RNA-seq data from various *Tetraselmis* spp. stains and calculate FPKM values for genes. The graphs show the boxplot distributions of FPKM values for genes sorted by VBM taxonomic categories and transcriptomes. FPKM values that differ significantly from the overall distribution in the category (i.e. outliers) are not shown. Two different sets of transcriptomes were analysed for the *T. striata* clade (**A—B)** and *T. suecica* clade (**C—E)**, as shown in the legends. The numbers of genes in each categories are given in Table S5.

Another question is why the measured levels of transcription are globally different between VBM categories. Phycodnaviridae best matching genes had systematically relatively high FPKM values (plus to some extent Mimiviridae, Pandoraviruses, and Pithoviridae) whereas TsV and TetV best matching genes were generally silent. Since most of TsV and TetV-like genes resided in genomic regions of viral origin, it appears that viral genes tend to be untranscribed. The origin of the other VBM gene categories is unknown (i.e., virus to alga HGT or the reverse) but the frequency of VBM genes that also show significant secondary BLASTP matches in *Viridiplantae* (i.e., includes green algae (*Chlorophyta*) and land plants (*Streptophyta*)) may give some indication in this regard. Indeed, secondary matches are consistent with the scenario that the gene is of algal origin but was recently captured by a virus, ultimately resulting in a best match in a virus derived from the lineage involved in the HGT. **Figure 6** shows that for each algal genome, the percentage of genes that have significant matches in Viridiplantae is higher for Phycodnaviridae, Mimiviridae, Pandoravirus, and Pithoviridae than for the other categories. This result therefore suggests that genes in these categories are more likely to have an algal rather than a viral origin. Taken together, these results lead to a consistent hypothesis that the categories Phycodnaviridae, Mimiviridae, Pandoravirus and Pithoviridae are characterized by higher transcriptional activity because they mainly consist of genes of algal origin that have been captured by viruses and whose expression is naturally required to participate in biological functions of the host cell.

**Figure 6.**
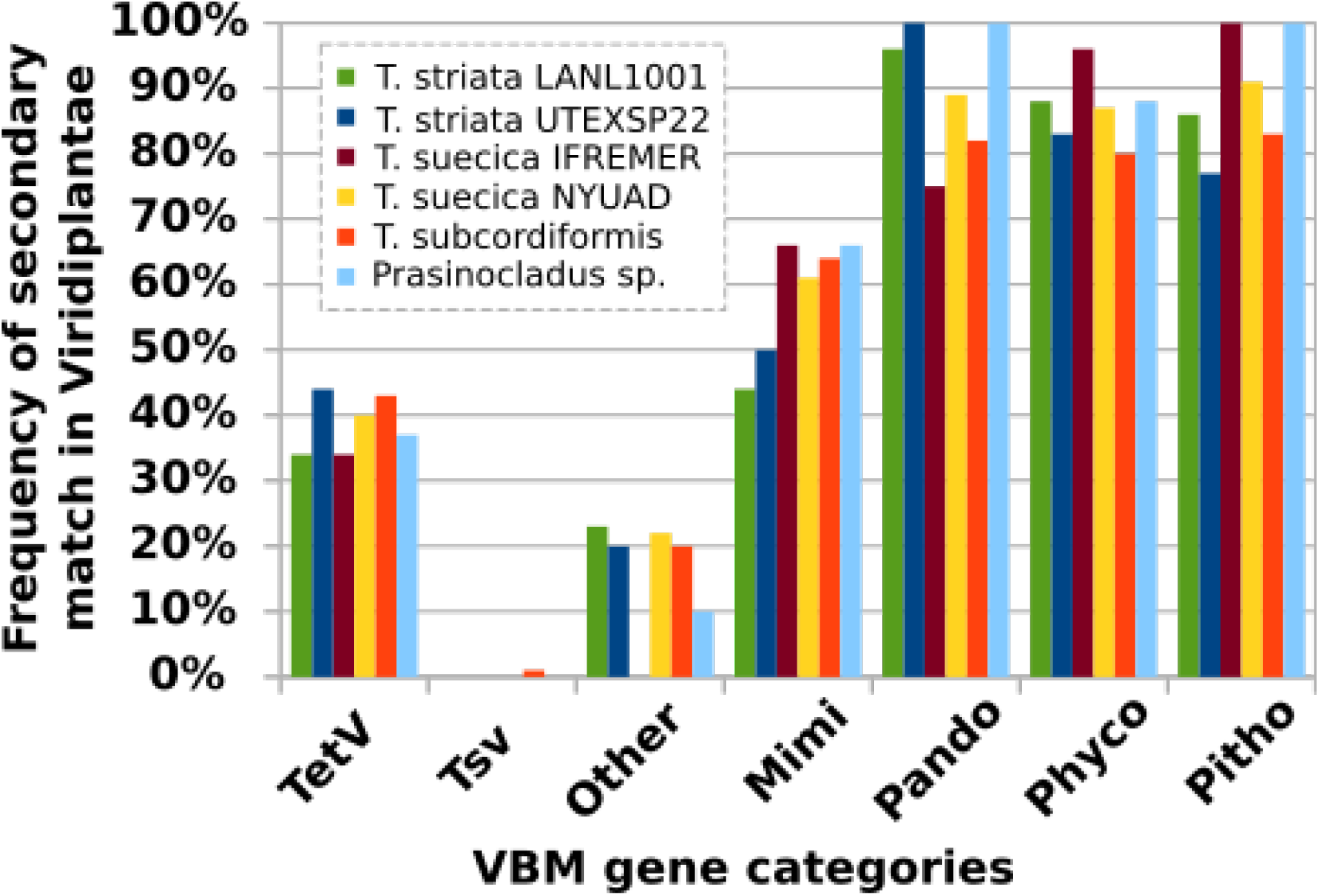
Percentage of genes with secondary matches in Viridiplantae.

## Discussion

### Transcriptionally silent giant virus-like insertion elements (phylum Nucleocytoviricota) in this study may be actually transcribed under specific conditions

Viral genomic regions in *T. striata* LANL1001 and *T. suecica* CCMP904 IFREMER that best match the algal giant virus TetV had GC content strikingly different than both the cellular host’s genome and the TetV genome. Positive windows containing TetV best matches in the other *Chlorodendraceae* genomes also had G+C content substantially higher than that of the TetV genome. Excluding the hypothesis that these GC content variations were the result of specific mutational bias within inserted viral regions induced by host DNA replication or DNA repair, it is highly likely that they reflect the diversity of GC contents that exists among TetV-related viruses.

Only a fraction of the TetV gene complement were found in the algal genomes. For instance neither of the two key fermentation genes (i.e., pyruvate formate-lyase and pyruvate formate-lyase activating enzyme), which have been a surprising discovery in the TetV genome (19), were found in the viral insertions or elsewhere in the algal genomes. Although we cannot rule out the viral genes were missed due to gaps in sequencing or because the viral genome insertions were incomplete, this result suggests that these accessory genes might have a sporadic distribution in the TetV-like clade, reflecting different adaptive niches.

In general, TetV-like genes found in algal genomes were not represented in *Chlorodendraceae* transcriptomes, suggesting that these genes do not produce mRNA, at least under the conditions under which the transcriptomes were generated. With the exception of LANL1001, it was not possible to align transcriptomes and genomes belonging to the same algal strain. Although overall, the results of the transcriptomic analyses were similar to those obtained for LANL1001, we cannot exclude that the lack of transcription measured in heterologous comparisons was the consequence of a physical absence of the corresponding genes in the algal strains used for transcriptome sequencing. This is especially true for *T. suecica* CCMP904 where a background low intensity infection of the stock culture is suspected. A complementary RNA-seq study of the CCMP904 culture would certainly be able to determine if this hypothesis is valid (viral genes are actively transcribed upon infection).

Giant virus genomic insertions are relatively common in algae and some protozoa but their biological role in the host cell remains unknown (13, 23, 44–46). This role could be nil if the viral DNA is permanently inactivated because the cellular transcription machinery is unable to recognize viral promoters or as the result of active gene silencing by the host. The absence of gene transcription in viral regions supports these hypotheses. Alternatively, viral-derived genes could be expressed under very specific conditions (e.g not unlike the response of genome integrated virophages to giant virus infection (8)), namely upon infection with a related virus that brings its transcription factors capable of recognizing viral promoters. Since viral host-borne genes have accumulated numerous mutations, their expression could lead to the production of abnormal viral proteins that would compete with the native viral proteins and hinder infection and/or the production of new infectious particles. Another possibility is that the inserted viral regions are transcribed into small interference RNAs (siRNAs), which are not covered by the transcriptomic sequencing data analyzed in this work. The siRNA pathway is an antiviral mecanism widespread across the eukaryotic phylogeny, which can potentially direct viral DNA methylation and transcriptional silencing (47). Obviously, further studies are needed to investigate and understand the biological significance of these giant-virus insertions.

### Polinton-like viruses likely exhibit dual lifestyles that also facilitate rapid hybridization

Chlorodendraceae genomes also showed an abundance of diverse PLV-like elements whose closest known relative was TsV infecting *T. striata*. TsV is the only PLV specimen that is known and isolated as a viral infectious particle. All other PLVs reported in the literature have been identified either as inserted elements in eukaryotic genomes or in viral metagenomes. For this reason, PLVs are suspected to have a dual lifestyle, alternating between a free viral phase and a phase integrated into host genomes. The PLV-like elements integrated into Chlorodendraceae genomes provide an additional argument to this hypothesis. It is highly likely that TsV is capable of inserting itself into the host genome under conditions that remain to be determined. Transcriptomic data, particularly in the case of LANL1001, suggest that PLV-like elements are essentially silent in their integrated form except for a gene homologous to TVSG_00024 of TsV that is transcribed in several elements. The function of this gene is unknown but given its quasi-systematic transcription in the inserted elements it is possible that the product of this gene is involved in a sensing mechanism that allows to trigger the reactivation/excision of the provirus according to a stimulus that remains to be discovered. In some transcriptomes, assembled sequences contained multiple ORFs encoding distinct PLV proteins. It is likely that these sequences correspond to distinct but overlapping transcription units (i.e., adjacent genes) that were assembled in the same contig. We do not know whether these transcripts were generated by integrated elements in the genomes of the concerned strains. However, given the very restricted transcription of the PLV elements of LANL1001, it is more likely that these highly transcribed genes result from an infection of the culture that served as a template for transcriptomic sequencing.

In the IFREMER strain, the distribution of elements in the genome was not random and was concentrated in a central region of the largest chromosome. It is possible that this region also houses the centromere, but this information is not known. In any case, this distribution suggests that insertions of PLVs occur in genomic hotspots. Further sequencing and near-complete assembly of the other algal strains will confirm whether these hotspots exist in other Chlorodendraceae. It is also striking that the Chlorodendraceae PLV elements had 3 different types of replication proteins (i.e., S3H helicase – TVpol fusion, pDNAP and S1H helicase) and that these were mutually exclusive, so that each element had only one gene. Given the relative phylogenetic proximity of PLV-elements to each other, this observation suggests that PLVs change their replication genes quite easily and frequently in evolution. The existence of element-dense integration hotspots, where new PLVs are likely to integrate into already established elements, is likely to favor the creation of hybrid elements that basically result in gene exchanges between PLVs, especially their replication gene. These new genetic combinations can then be tested by natural selection if the hybrid genome can be excised and propagated in a neoformed virion. In this context, it is interesting to note the existence of hybrids between virophages and polintons, as was found among rumen virophages where a virophage MCP was combined with a polinton DNA polymerase B (48)

## Conclusion

This study explored both genomic and transcriptomic data of previously sequenced algae (family *Chlorodendraceae*) with close relation to a polinton-like virus (TsV) host *T. striata* to further support the dual lifestyle hypothesis among PLVs. Additionally, integration elements of giant viruses were also explored, given a known *Tetraselmis* sp. virus (TetV). Both of these analyses uncovered integrated viral elements among several of the related algal species and gave insights into their behaviors. Given the substantial evidence of both giant virus and PLV viral integrated elements, we recommend future studies target the mechanism(s) behind triggering the possible transcription of these integration elements. More specifically we recommend further characterization of the unknown protein, TVSG_00024 (found within many PLV integrated viral elements) given its possible role in triggering transcription of PLV viral integrated elements. This work could further reveal the mechanisms behind the implied dual lifestyle of PLVs.

## Abbreviations

MCP: major capsid protein
mCP: minor capsid protein
PLV: polintons- like virus
TsV: T. striata virus
PgVV: Phaeocystis globasa virus virophage
ORF: open reading frame
TIRs: terminal inverted repeats

## Acknowledgment

We would like to thank Rossana Sussarellu, Gregory Carrier and Sabine Stachowski- Haberkorn from IFREMER Nantes for providing us access to their CCMP904 data in advance of publication.

