## Supplementary material for "Integrated viral elements suggest the dual lifestyle of *Tetraselmis* spp. polinton-like viruses": SUPP: Fig S1.pdf

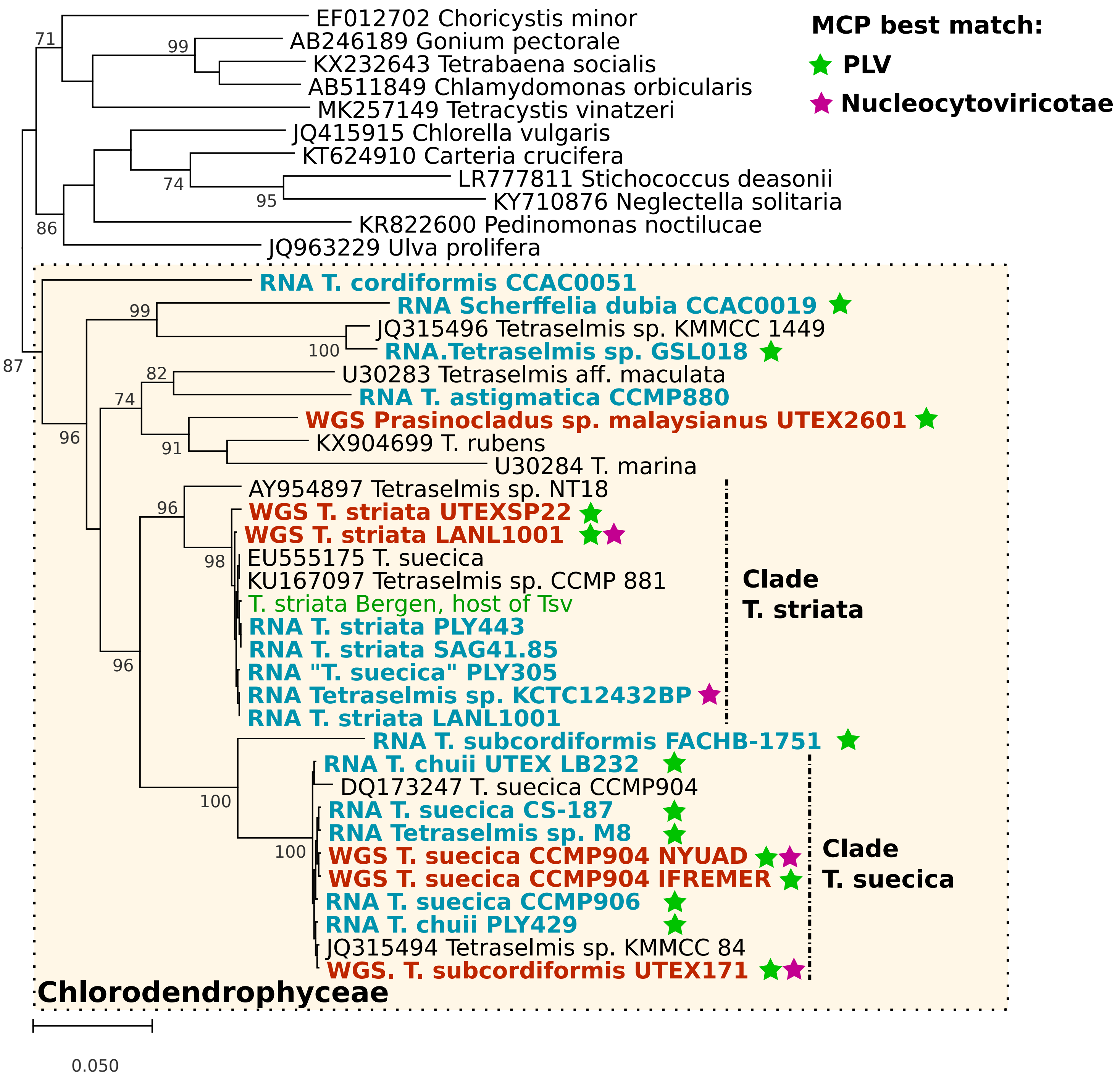


Figure S1: Phylogenetic reconstruction of Chlorodendrophyceae based on the RBCL coding sequence (nucleotides). Sequences from genomic assemblies are shown in red, while sequences from assembled transcriptomes are shown in blue. The 18S gene sequence of T. striata "Bergen", i.e., the strain from which Tsv was first isolated, was determined in this study (sequence available in the multiple alignment given as supplementary data). The other sequences were downloaded from Genbank and are shown in black with their accession number. The green and purple stars indicate datasets that contained MCP sequences with a best match against PLV or Nucleocytoviricota, respectively (as listed in Table 1).
