## Supplementary material for "Integrated viral elements suggest the dual lifestyle of *Tetraselmis* spp. polinton-like viruses": SUPP: Fig S2.pdf

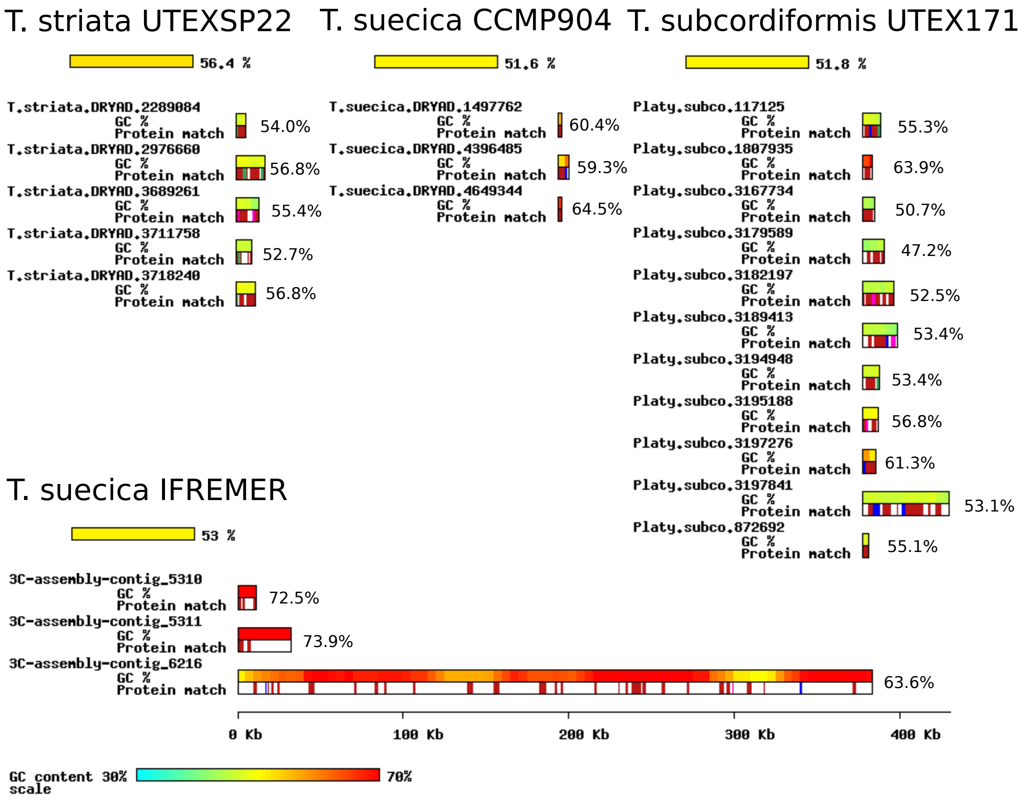


Figure 4. Viral genomic regions identified in the algal genome assemblies. Representation of GC content and ORF best matches are the same as in Figure 2. The average GC content of the alga genomes is provided below the species name.
