## Supplementary material for "Integrated viral elements suggest the dual lifestyle of *Tetraselmis* spp. polinton-like viruses": SUPP: Fig S3.pdf

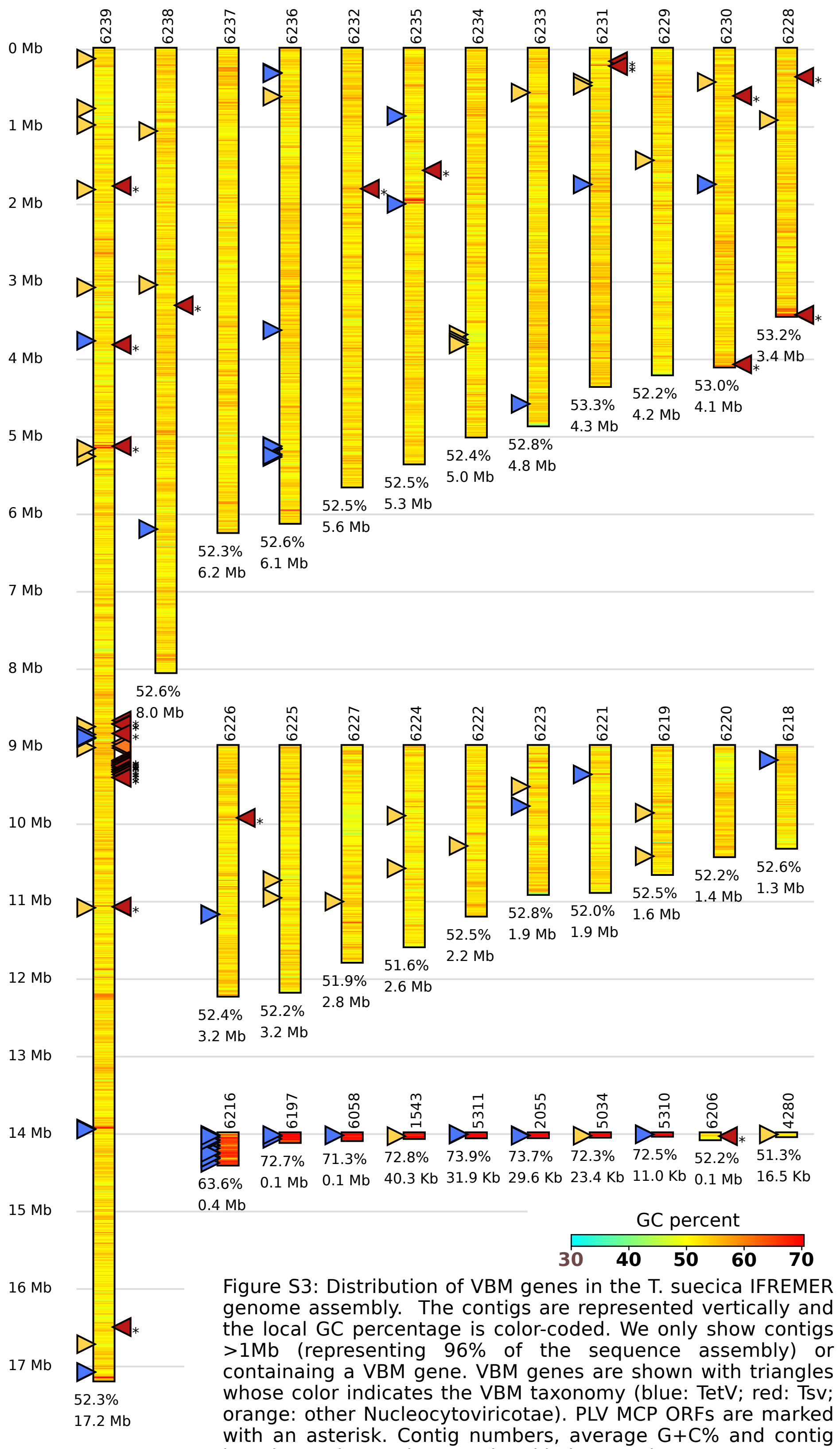


Figure S3: Distribution of VBM genes in the T. suecica IFREMER genome assembly. The contigs are represented vertically and the local GC percentage is color-coded. We only show contigs >1Mb (representing 96% of the sequence assembly) or containaing a VBM gene. VBM genes are shown with triangles whose color indicates the VBM taxonomy (blue: TetV; red: Tsv; orange: other Nucleocytoviricotae). PLV MCP ORFs are marked with an asterisk. Contig numbers, average G+C% and contig length are shown above and and below contigs.
