## Supplementary material for "Integrated viral elements suggest the dual lifestyle of *Tetraselmis* spp. polinton-like viruses": SUPP: Fig S4.pdf

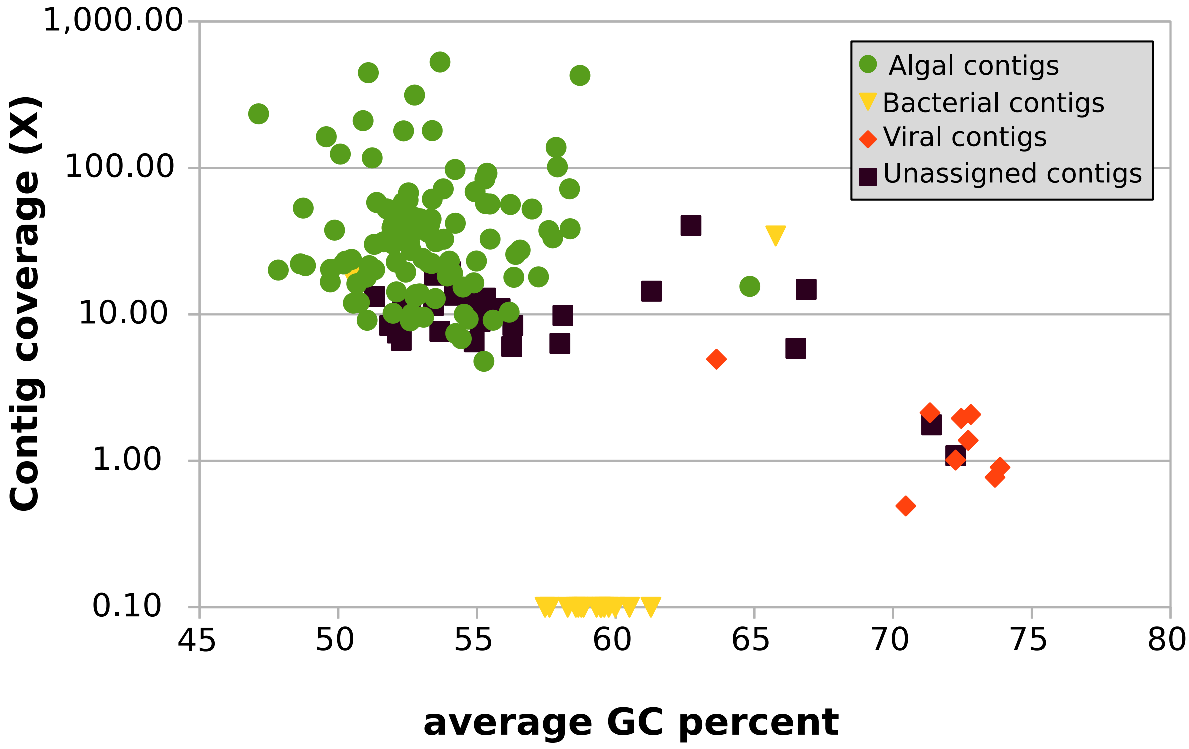


Figure S4: Coverage depth of IFREMER CCMP904 contigs mapped with NYUAD illumina reads. Contigs are represented by symbols whose color reflects the taxonomic assignation as shown in the Figure. The vast majority of bacterial contigs had zero coverage but were assigned a value of X=0.1 to be included in the Figure (y-axis in logarithmic scale). These contigs represent specific bacterial contamination of the IFREMER culture.
