## Supplementary material for "Integrated viral elements suggest the dual lifestyle of *Tetraselmis* spp. polinton-like viruses": SUPP: Fig S5.pdf

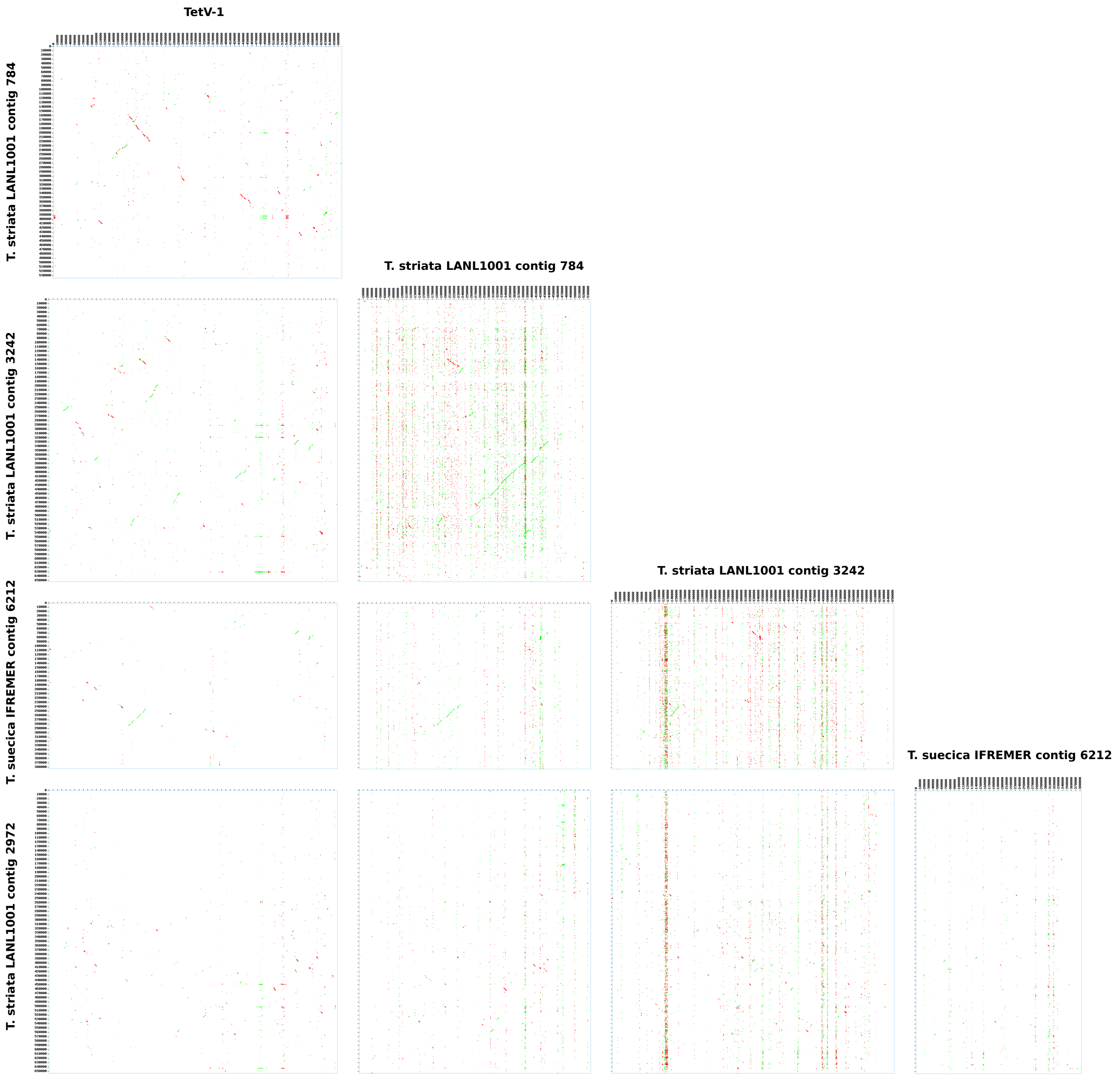


Figure S5: dot plot alignments of selected algal contigs and the TetV-1 genome

Red and gree dots represent significant TBLASTX matches (evalue<0.001) in forward of reverse directions, respectively
