## Supplementary material for "Integrated viral elements suggest the dual lifestyle of *Tetraselmis* spp. polinton-like viruses": SUPP: Fig S6 A&B.pdf

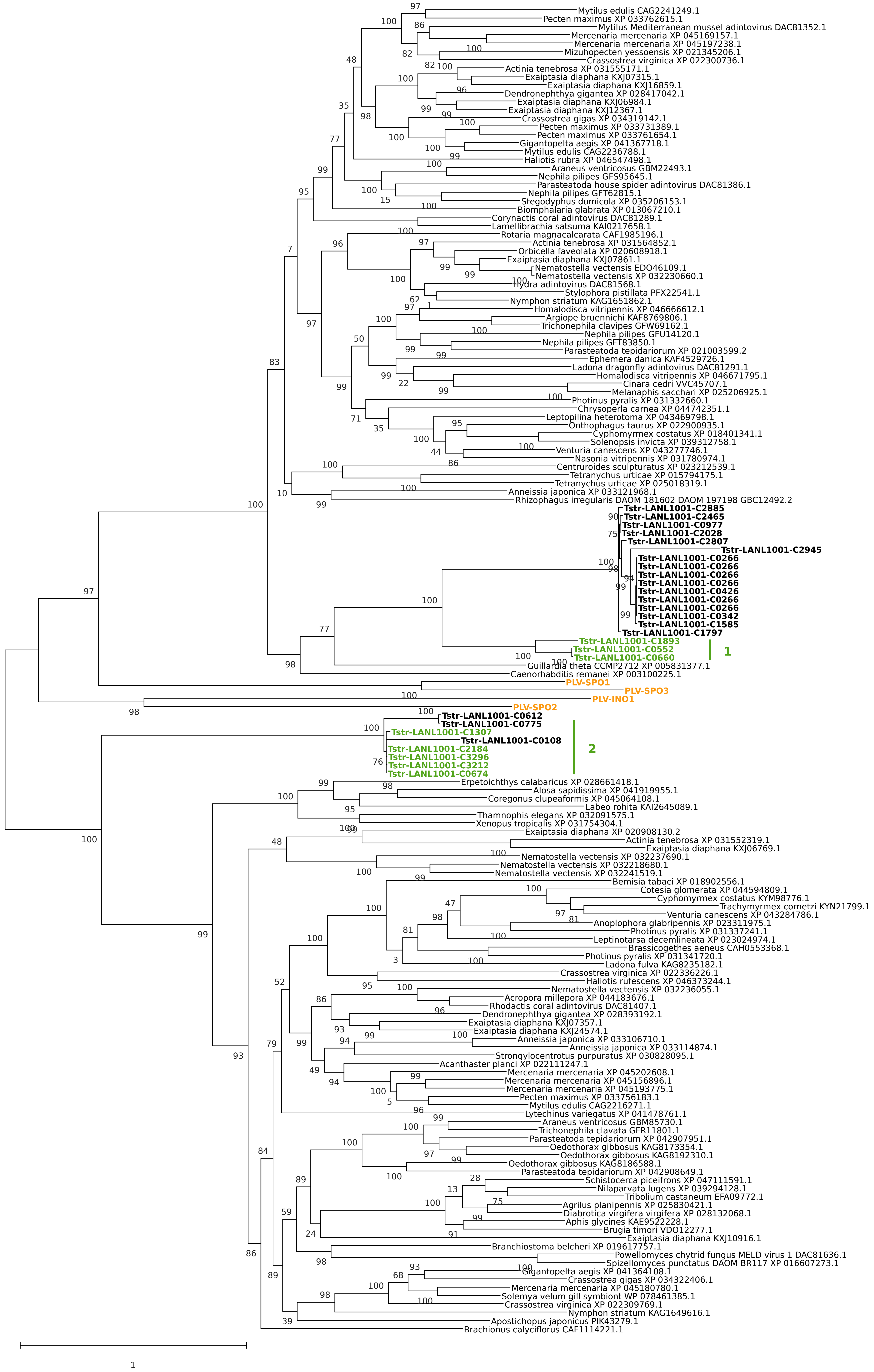


Figure S6A. Phylogenetic reconstruction of the PLV pDNAP proteins. In green, the proteins encoded by PLV elements shown in Fig. 4. Two phylogenetic groups of PLV pDNAP, 1 and 2, are delineated in green. In black and bold are Chlorodendraceae proteins that were not idenfied as belonging to a PLV element (i.e., not in the vicinity of a MCP protein).

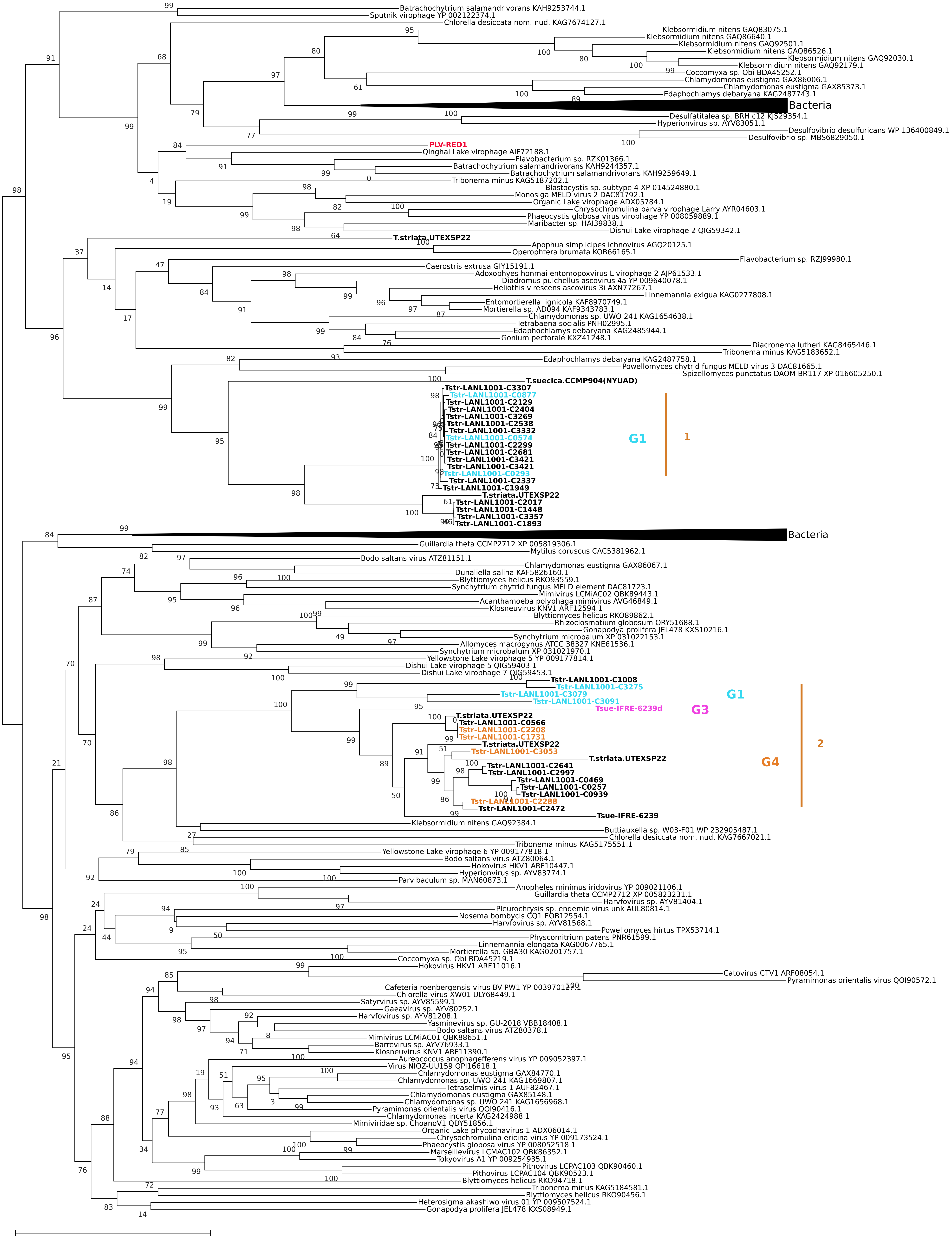


Figure S6B. Phylogenetic reconstruction of the S3H helicase – TVpol fusion proteins. Proteins encoded by PLV elements shown in Fig. 4 are color coded. In blue, mauve and orange indicate proteins from G1, G3 and G4 element groups defined in Fig. 4, respectively. In black and bold are Chlorodendraceae proteins that were not idenfied as belonging to a PLV element (i.e., not in the vicinity of a MCP protein). Two phylogenetic groups of PLV S3H-TVpol proteins, 1 and 2, are delineated in orange (information reported on Fig. 4).
