## Supplementary material for "Integrated viral elements suggest the dual lifestyle of *Tetraselmis* spp. polinton-like viruses": SUPP: Fig S7.pdf

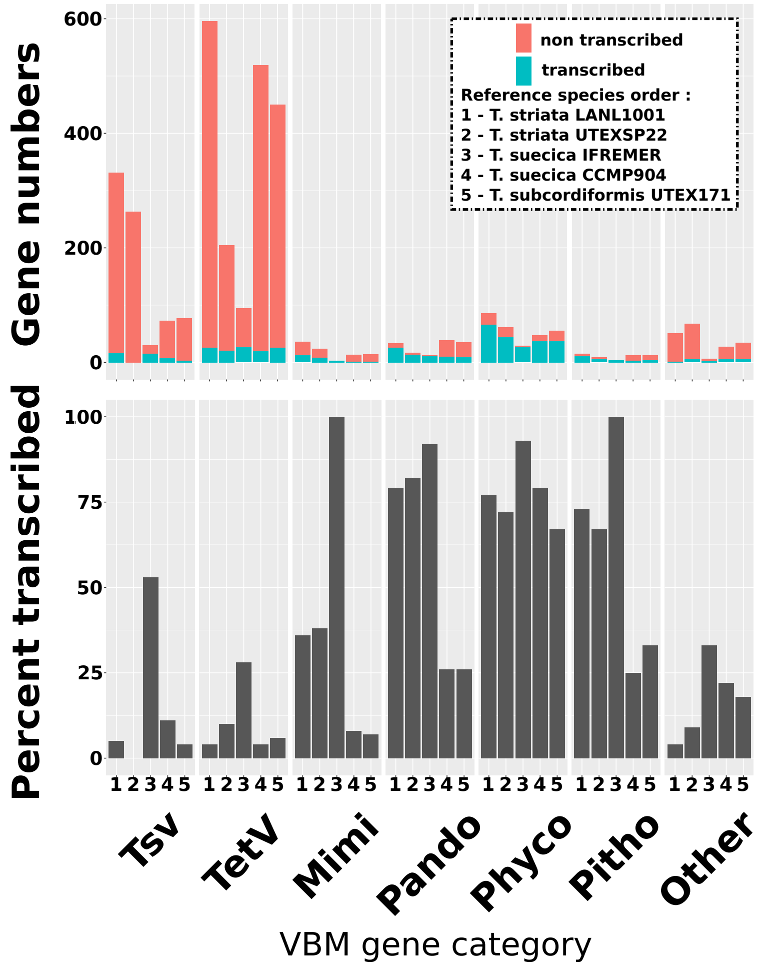


Figure S7: Transcription of VBM genes
(A) Number of transcribed versus non transcribed VBM genes of reference genome assemblies classified in taxonomic categories. A gene was considered as transcribed when its FPKM value was >1 in at least one of the aligned Chlorodendraceae transcriptomes (see Figure 5). (B) percent of transcribed VBM genes in the category.
